# Next-generation genetically encoded biosensors for spatiotemporal intracellular pH monitoring, mapping, and profiling

**DOI:** 10.64898/2026.05.19.726343

**Authors:** Sam Taylor, Bruno Colon, Kyutae D. Lee, Jennifer Arcuri, Shraddha Chandthakuri, Daniel G. Isom

**Affiliations:** Department of Molecular and Cellular Pharmacology, University of Miami Miller School of Medicine, Miami, FL, 33136, USA; Sylvester Comprehensive Cancer Center, University of Miami Miller School of Medicine, Miami, FL, 33136, USA; Frost Institute for Data Science and Computing, Coral Gables, FL, USA; University of Miami Medical Scientist Training Program, University of Miami Miller School of Medicine, Miami, FL 33136, USA; The Sheila and David Fuente Graduate Program in Cancer Biology; Molecular and Cellular Pharmacology Graduate Program

## Abstract

Bioluminescence resonance energy transfer (BRET) systems are widely used for live-cell spectroscopy and biosensor engineering, yet the intrinsic pH sensitivity of commonly used BRET components has not been systematically examined. Here, we show that major BRET luciferase donors, fluorescent acceptors, and donor–acceptor assay pairs exhibit pronounced pH-dependent spectroscopic behavior across physiologically relevant conditions, identifying environmental pH responsiveness as a fundamental property of widely used BRET systems and a potential source of previously underappreciated assay artifacts. Leveraging these principles, we engineered ORION (ratiOmetRIc prOton seNsor), a genetically encoded ratiometric BRET pH sensor based on the NanoLuc–mVenus fusion. ORION exhibited strong brightness, an approximately 9-fold dynamic range, and robust responsiveness across a substantially broader pH range than that of existing genetically encoded sensors. Compared to pHluorin2, ORION maintained substantially improved quantitative performance at acidic pH values below 6.0. To demonstrate its utility in a biological application, we applied ORION across diverse cancer cell models and identified heterogeneous acid imprinting states, suggesting that tumor cells can retain persistent physiological memory of adaptation to acidic microenvironments even after prolonged ex vivo culture. Together, these findings establish pH responsiveness as a fundamental property of BRET systems and position ORION as a best-in-class platform for interrogating and quantifying pH regulation of biology in living systems.

## Introduction

Proton concentration is a fundamental regulator of biological systems across molecular, cellular, and tissue scales. pH influences protein conformation^1^, enzymatic activity^2^, membrane transport^3^, metabolism^4,5^, signaling^6,7 1,8,9^, and organelle identity ^10^, while spatially resolved pH gradients help define intracellular compartments, including endosomes, lysosomes, and secretory vesicles^11,12^. Dynamic regulation of pH is also central to organismal physiology and disease. In cancer, extracellular acidification promotes invasion, immune evasion, metabolic adaptation, and therapeutic resistance^4,13^, while altered pH homeostasis has similarly been implicated in inflammation, ischemia, infection, and neurodegeneration^10,14-16^. Because pH is highly dynamic and heterogeneous across biological environments, technologies capable of accurately measuring pH in living systems are essential for understanding both normal physiology and disease-associated processes.

Optical approaches have become the dominant strategy for measuring biological pH dynamics in living systems. These approaches broadly include small-molecule fluorescent dyes and genetically encoded sensors. Widely used fluorescent dyes, including BCECF^14^, SNARF^17^, and LysoSensor^14^ derivatives, have enabled measurements of cytosolic, extracellular, and organellar pH across diverse biological contexts. In parallel, genetically encoded sensors such as pHluorin^18,19^, pHuji^20^, pHRed^21^, and related protein- and nucleic acid-based architectures^14,22-24^ have expanded the ability to monitor pH dynamics, with subcellular targeting and longitudinal imaging capabilities. Despite their broad utility, fluorescence-based approaches remain associated with important limitations, including photobleaching, excitation-induced autofluorescence, phototoxicity, spectral overlap, and reduced performance in highly acidic environments. These challenges have motivated growing interest in luminescence-based sensing strategies that avoid external excitation while maintaining high sensitivity in living systems^25-27^.

Bioluminescence resonance energy transfer (BRET) has emerged as a powerful spectroscopic platform for studying biological processes in living cells and organisms. In BRET systems, energy generated by a bioluminescent donor luciferase is transferred non-radiatively to an acceptor fluorophore, producing a ratiometric optical output that is sensitive to donor-acceptor interactions and local molecular environments^26,28^. Because BRET does not require excitation light, these systems exhibit low background signal, reduced phototoxicity, and improved compatibility with deep-tissue and longitudinal imaging applications. As a result, BRET-based technologies have been widely adopted for applications including G protein-coupled receptor (GPCR) signaling assays^26,29-32^,protein-protein interaction studies, trafficking measurements, biosensor engineering, and drug discovery. Collectively, these features position BRET systems as versatile and highly sensitive platforms for live-cell spectroscopy.

During our studies involving BRET-based GPCR assays, we observed substantial signal changes following extracellular acidification. Because extracellular pH changes drive intracellular acidification in mammalian cells^7^, we hypothesized that commonly used BRET donors and acceptors might themselves exhibit intrinsic pH sensitivity. This possibility raised two important implications. First, pH-dependent changes in donor and acceptor behavior could introduce previously underappreciated artifacts into the interpretation of BRET-based assays. Second, the intrinsic environmental responsiveness of BRET systems might be rationally leveraged to engineer new genetically encoded pH biosensors. Surprisingly, despite the widespread adoption of BRET technologies across biological research, systematic characterization of pH sensitivity across commonly used BRET components and assay architectures has not been reported.

Here, we systematically characterize the pH sensitivity of widely used BRET donors, acceptors, and assay pairs across physiologically relevant conditions. We find that major BRET components exhibit pronounced, tunable pH responsiveness, demonstrating pH sensitivity as a fundamental property of commonly used BRET systems. Leveraging these principles, we engineer ORION (ratiOmetRIc prOton seNsor), a genetically encoded ratiometric BRET pH sensor optimized for live-cell imaging and translational applications. We benchmark ORION against established fluorescent pH sensors, including pHluorin, and demonstrate enhanced performance across acidic pH ranges relevant to biological and disease-associated environments. Finally, we apply ORION to characterize acid imprinting across multiple cancer cell models, establishing its utility for interrogating pH dynamics in living systems.

## Results

### Common BRET components are highly sensitive to pH

To systematically characterize the pH responsiveness of commonly used BRET components, we leveraged *Saccharomyces cerevisiae* (hereafter, yeast) as an experimentally tractable screening system. Yeast enables straightforward plasmid-based expression and direct spectroscopic measurements in intact cells without the need for protein purification^1,6-9,33,34^. In addition, mild permeabilization permits controlled equilibration of intracellular pH conditions while preserving measurements within a cellular context. Collectively, these features provided a practical framework for benchmarking donor and acceptor pH responsiveness across physiologically relevant conditions and pH values.

Using this system, we examined the pH responsiveness of the luciferase donors NanoLuc (Nluc) and Renilla luciferase 8 (Rluc8), together with the fluorescent acceptors GFP2 and mVenus (Fig. 1a,b). Following plasmid-based expression in yeast, cells were mildly permeabilized with digitonin to permit equilibration of the cytoplasm with exogenous buffers spanning pH 5.0–8.5 prior to spectroscopic measurements. This approach enabled direct comparison of donor and acceptor behavior across a wide range of pH values commonly encountered in biological systems and frequently modeled in experimental studies.

**Figure 1.**
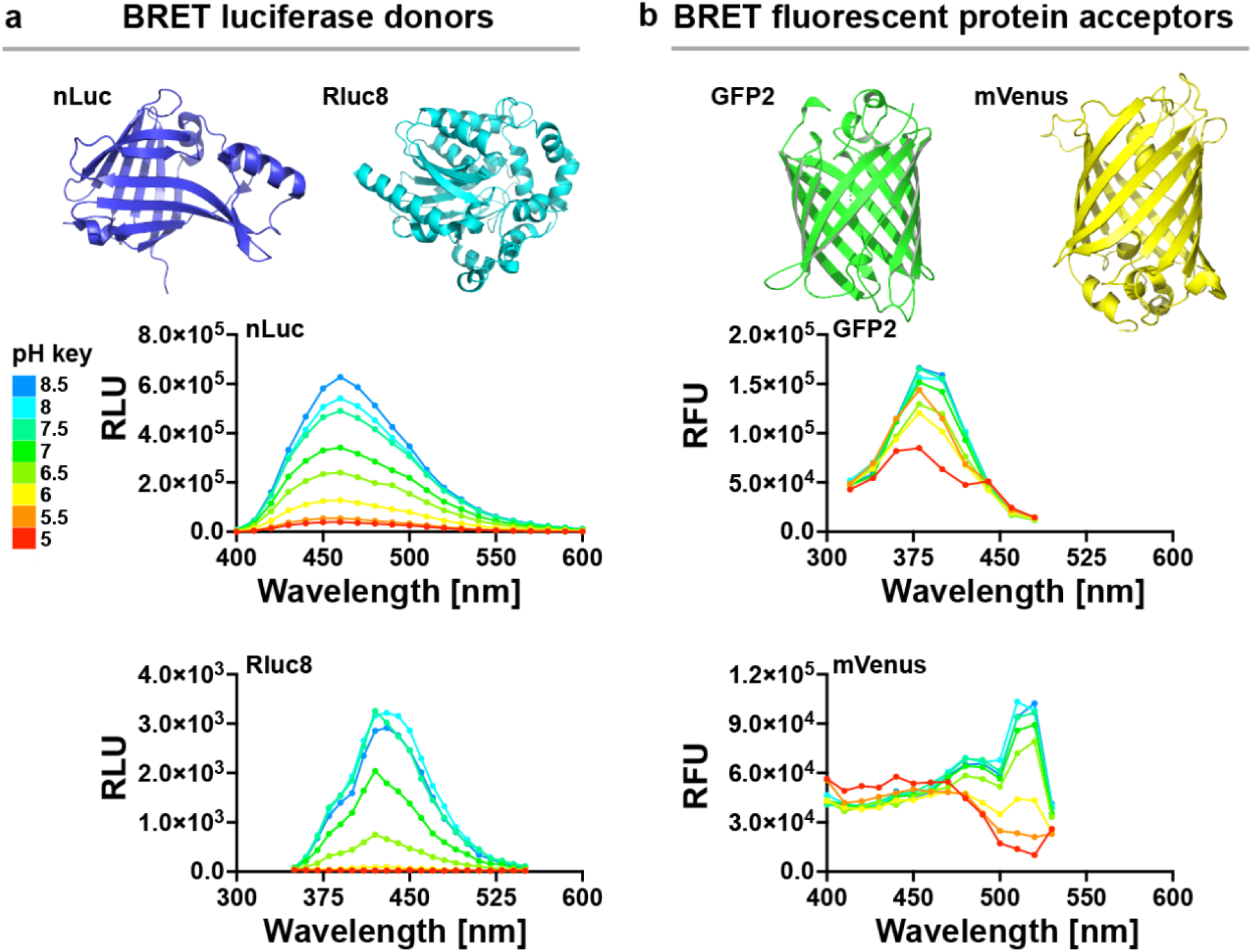
pH sensitivity of commonly used BRET donor and acceptor components. (**a**) Experimentally resolved structures of the widely used BRET luciferase donors NanoLuc (Nluc; PDB: 7SNS) and Renilla luciferase 8 (Rluc8; PDB: 6YN2). Bioluminescence emission spectra of Nluc and Rluc8 were measured across pH 5.0–8.5 in digitonin-permeabilized *S. cerevisiae* following addition of the luciferase substrate furimazine. Both luciferase donors exhibited pronounced pH-dependent changes in emission intensity across physiologically relevant conditions. (**b**) Experimentally resolved structures of the widely used BRET fluorescent protein acceptors GFP2 (PDB: 1EMC) and mVenus (PDB: 7PNN). Fluorescence excitation spectra of GFP2 and mVenus were measured across pH 5.0–8.5 under identical conditions. Both fluorescent acceptors displayed substantial pH-dependent spectral responses, demonstrating that pH sensitivity is a shared property of commonly used BRET donor and acceptor components. Data represent mean values from n = 3 independent experiments.

Both luciferase donors displayed pronounced pH-dependent changes in emission intensity (Fig. 1a). NanoLuc showed particularly strong responsiveness, with maximal luminescence at alkaline pH and progressive attenuation as pH decreased. Renilla luciferase 8 similarly demonstrated substantial pH sensitivity, although with a distinct spectral profile and lower overall emission intensity. These findings indicate that luciferase donors commonly used in BRET assays are not spectroscopically invariant across physiological pH ranges, but instead exhibit substantial environmental responsiveness that can influence assay output.

We next examined whether fluorescent acceptor proteins paired with these luciferase donors similarly exhibit intrinsic pH sensitivity. GFP2 and mVenus each displayed marked pH-dependent excitation responses across the tested range (Fig. 1b). GFP2 showed progressively reduced excitation efficiency under acidic conditions, whereas mVenus exhibited pronounced spectral alterations and signal suppression at lower pH values. Together, these results demonstrate that pH responsiveness is a broader property shared by both donor and acceptor components in BRET systems, suggesting that pH fluctuations can substantially influence BRET assay behavior while also providing an opportunity to exploit these intrinsic spectroscopic properties for pH biosensor engineering.

### Common BRET pairs are highly sensitive to pH

Having established that individual donor and acceptor components exhibit substantial intrinsic pH responsiveness, we next examined how these effects manifest within intact BRET donor–acceptor architectures. To address this, we generated fusion constructs pairing the luciferase donors NanoLuc (nLuc) or Renilla luciferase 8 (Rluc8) with the fluorescent acceptors mVenus or GFP2 through flexible GGSGGS linkers (Fig. 2a,b). AlphaFold2 structural modeling suggested that these constructs position donor and acceptor chromophores within distances compatible with efficient resonance energy transfer while maintaining substantial conformational flexibility. We therefore reasoned that pH-dependent effects on either donor emission or acceptor excitation could become amplified within the context of coupled BRET systems.

**Figure 2.**
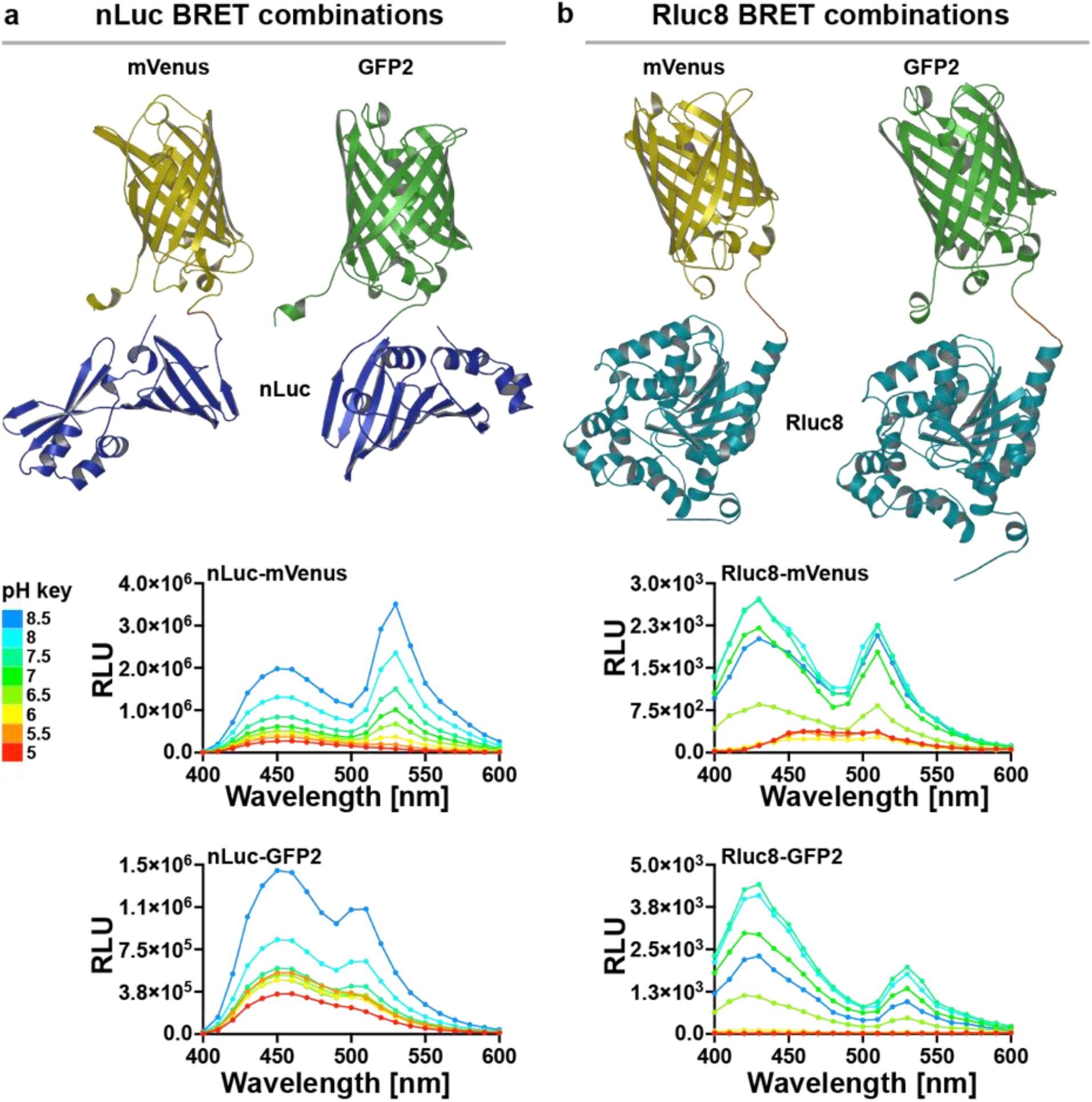
Intrinsic pH sensitivity of commonly used BRET donor–acceptor assay pairs. (**a**) AlphaFold2-predicted structures of NanoLuc (nLuc) fused to the fluorescent protein acceptors mVenus and GFP2 through flexible GGSGGS linkers, shown in orange. Bioluminescence emission spectra of nLuc–mVenus and nLuc–GFP2 fusion constructs were measured across pH 5.0–8.5 in digitonin-permeabilized *S. cerevisiae* following addition of the luciferase substrate furimazine. NanoBRET pairs exhibited pronounced pH-dependent changes in both donor and acceptor emission profiles across physiologically relevant conditions. (**b**) AlphaFold2-predicted structures of Renilla luciferase 8 (Rluc8) fused to the fluorescent protein acceptors mVenus and GFP2 through flexible GGSGGS linkers, shown in orange. Bioluminescence emission spectra of Rluc8–mVenus and Rluc8–GFP2 fusion constructs were measured across pH 5.0–8.5 under identical conditions. Renilla BRET assay pairs similarly displayed substantial pH-dependent changes in emission intensity and spectral output, demonstrating that intrinsic pH responsiveness is a conserved property across major BRET architectures. Data represent mean values from n = 3 independent experiments.

Using the same digitonin-permeabilized yeast platform described above, we measured full bioluminescence emission spectra for each BRET pair across buffers spanning pH 5.0–8.5 following substrate addition. All four donor–acceptor combinations displayed pronounced pH-dependent alterations in spectral output (Fig. 2a,b). In each case, acidification progressively suppressed overall signal intensity while also reshaping the relative balance between donor and acceptor emission peaks. These effects demonstrate that commonly used BRET architectures are not spectroscopically stable across physiological pH ranges, but instead integrate the intrinsic environmental sensitivities of both donor and acceptor components into complex pH-responsive behaviors.

Among the NanoLuc-based architectures, the nLuc–mVenus pair exhibited the largest and most coordinated pH-dependent spectral shifts (Fig. 2a). Alkaline conditions produced strong acceptor emission centered near 530 nm, accompanied by robust overall signal intensity, whereas acidic conditions markedly suppressed acceptor output and reduced total luminescence. Importantly, these transitions occurred smoothly across the tested pH range, producing large dynamic spectral changes well suited for ratiometric quantification. In contrast, the nLuc–GFP2 pair showed comparatively weaker acceptor emission and a more compressed dynamic range under acidic conditions.

The Rluc8-based architectures similarly exhibited substantial pH responsiveness, although with lower overall emission intensity and reduced spectral separation between donor and acceptor peaks (Fig. 2b). Both Rluc8–mVenus and Rluc8–GFP2 demonstrated progressive attenuation under acidic conditions, but their lower brightness and diminished acceptor contrast suggested reduced suitability for sensitive live-cell measurements relative to NanoLuc-based systems.

Collectively, these results demonstrate that pH sensitivity is an emergent property of intact BRET architectures rather than an isolated feature of individual donor or acceptor proteins. Among the combinations tested, the nLuc–mVenus pair exhibited the strongest brightness, the largest ratiometric spectral shifts, and the broadest dynamic responsiveness across physiologically relevant pH conditions, motivating its selection for development as a genetically encoded intracellular pH biosensor termed ORION.

### Characterization, benchmarking, and validation of the ORION pH sensor

To quantitatively define the sensing properties of ORION independent of cellular context, we next purified recombinant sensor components and systematically characterized their spectroscopic behavior across pH 4.0–8.5. We further benchmarked ORION against pHluorin2, a widely used genetically encoded fluorescent pH sensor, to compare dynamic range, operational pH sensitivity, and performance under acidic conditions relevant to tumor biology and intracellular organelles.

We first examined the intrinsic pH responsiveness of the isolated ORION donor and acceptor components. Purified NanoLuc displayed strong pH-dependent changes in emission intensity across pH 4.0–8.5, with maximal luminescence under alkaline conditions and progressive attenuation at lower pH values (Fig. 3a). Purified mVenus similarly exhibited pronounced pH-dependent excitation behavior, including marked suppression under acidic conditions. These findings demonstrated that both components independently contribute substantial environmental responsiveness across physiologically and pathologically relevant pH ranges, suggesting that coupling these spectroscopic behaviors within a BRET architecture could generate amplified ratiometric sensing properties.

**Figure 3.**
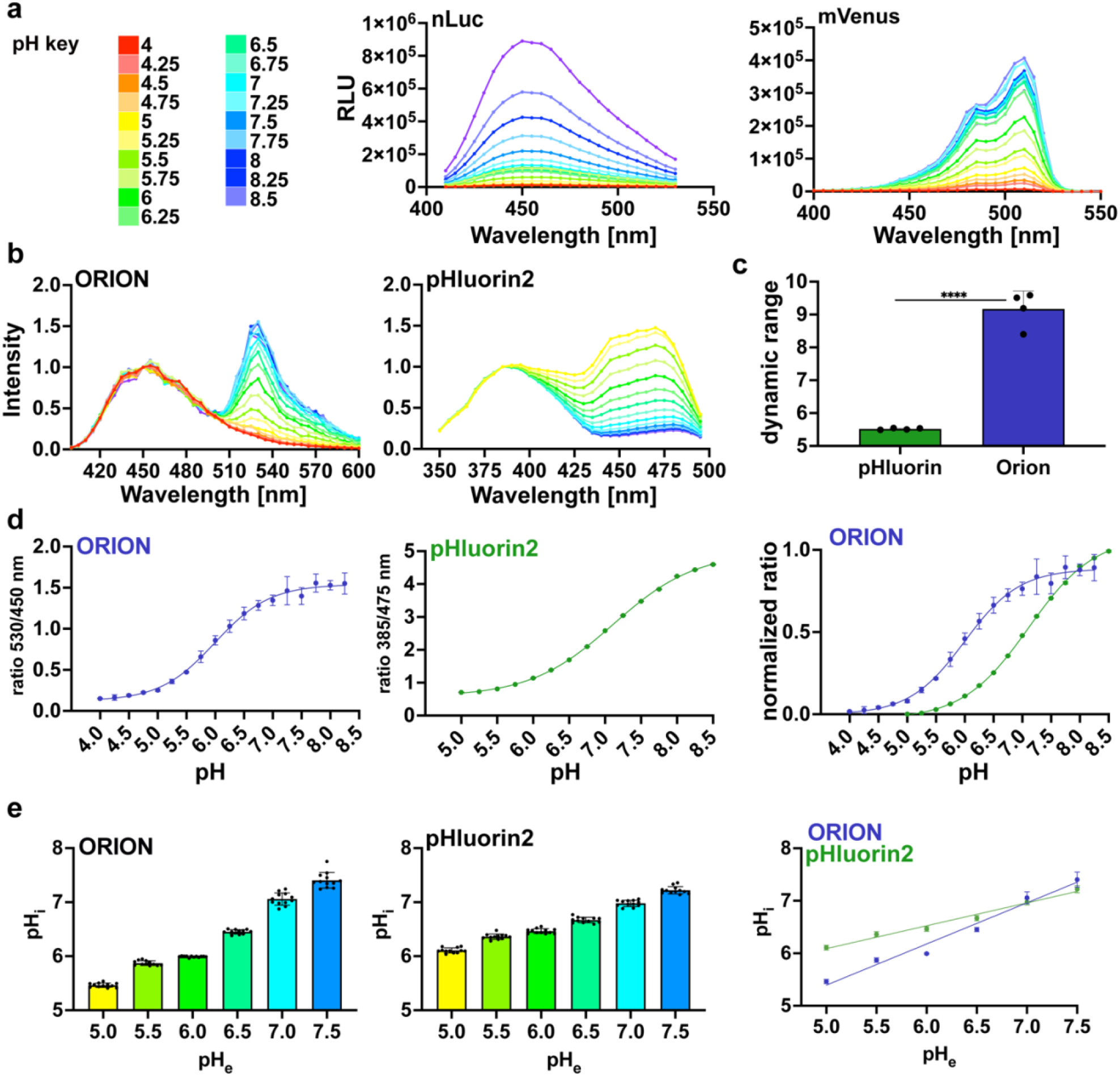
Characterization, benchmarking, and validation of the ORION pH sensor. Unless otherwise indicated, spectroscopic measurements were performed using purified recombinant proteins expressed in *E. coli*. (**a**) pH color key and emission spectra of purified NanoLuc (nLuc; left) and fluorescence excitation spectra of purified mVenus (right) measured across pH 4.0–8.5, demonstrating pronounced intrinsic pH-dependent spectral responses in the isolated donor and acceptor components used to engineer ORION. (**b**) Full BRET emission spectra of purified ORION (left) and fluorescence excitation spectra of purified pHluorin2 (right) measured across pH 4.0–8.5 and normalized to the lower wavelength peak of each sensor. ORION exhibited strong ratiometric spectral separation across acidic and physiological pH ranges. (**c**) Dynamic range comparison between ORION and pHluorin2, calculated as the fold-change between maximal and minimal ratio values across the measured pH range. ORION displayed a significantly larger dynamic range than pHluorin2 (mean ± SD; ****P < 0.0001; unpaired two-tailed Student’s *t*-test; n = 4 independent experiments). (**d**) ORION (530/450 nm; left) and pHluorin2 (385/475 nm; center) ratiometric pH titration curves across pH 4.0–8.5 fit using four-parameter logistic (4PL) regression models. A normalized overlay of both titration curves (right) demonstrates that ORION maintains substantial dynamic responsiveness across acidic pH conditions, where pHluorin2 sensitivity is reduced. (**e**) Intracellular pH (pH_i_) measurements as a function of extracellular pH (pH_e_) in HEK293T cells expressing ORION (left) or pHluorin2 (center) across extracellular pH 5.0–7.5. Overlay of both sensors (right) demonstrates concordant intracellular pH measurements across overlapping ranges, while revealing that ORION retains robust responsiveness under acidic conditions below pH 6.0, where pHluorin2 sensitivity is compressed. Data represent mean ± SD from n = 3 biological replicates with 4 technical replicates per condition.

We next measured full emission spectra of purified ORION across the same pH range and compared its behavior to pHluorin2 (Fig. 3b). ORION exhibited large and continuous ratiometric spectral transitions spanning acidic to physiological pH conditions. Acidification progressively reduced acceptor emission near 530 nm relative to donor emission near 450 nm, generating robust wavelength redistribution across the measured range. In contrast, pHluorin2 displayed comparatively compressed spectral changes under acidic conditions, with much of its dynamic responsiveness occurring closer to neutral and alkaline pH values.

Quantitative benchmarking demonstrated that ORION substantially outperformed pHluorin2 in overall dynamic range (Fig. 3c). Across the tested pH interval, ORION exhibited approximately a nine-fold ratiometric response window, whereas pHluorin2 showed a substantially narrower response range. Importantly, ORION maintained strong responsiveness under acidic conditions, which are frequently encountered in tumor microenvironments, endolysosomal compartments, and metabolically stressed tissues.

Ratiometric titration analysis further revealed distinct operational regimes for the two sensors (Fig. 3d). ORION displayed a broad, smoothly distributed transition spanning acidic to physiological pH values, whereas pHluorin2 showed a right-shift toward more alkaline conditions. Consistent with its lower apparent pKa (∼6.0 versus ∼7.0 for pHluorin2), ORION maintained strong responsiveness across a wider pH range (4.5–8.0), while the usable range of pHluorin2 was compressed. Combined with its substantially larger dynamic range, these properties make ORION ideal for quantifying the full spectrum of biological pH environments encountered in mammalian systems, including acidic compartments and microenvironments that are difficult to resolve using existing genetically encoded sensors.

Lastly, to validate ORION in living cells, we leveraged our established observation that acute extracellular acidification drives corresponding reductions in intracellular pH (pHi) in mammalian cell culture systems ^1,7^. HEK293T cells expressing either ORION or pHluorin2 were equilibrated in extracellular buffers spanning pH 5.0–7.5, and intracellular pH measurements were quantified from the resulting sensor responses (Fig. 3e). Both sensors produced concordant intracellular pH measurements across overlapping physiological ranges, supporting the quantitative accuracy of ORION in intact cells.

However, under strongly acidic extracellular conditions, the measurements progressively diverged as pHluorin2 responsiveness became increasingly compressed, consistent with reduced quantitative accuracy near the lower limit of its operational range. In contrast, ORION retained robust dynamic responsiveness across acidic conditions whereas pHluorin2 sensitivity diminished substantially. These results validate ORION’s ability to accurately quantify intracellular acidification across physiologically and pathologically relevant ranges. Together, these findings establish ORION as a sensitive genetically encoded ratiometric pH biosensor with robust performance across acidic and physiological pH environments relevant to cancer biology, vesicular trafficking, and cellular stress responses.

### Characterizing acid imprinting across cancer cell models

Having established the sensitivity and quantitative performance of ORION, we next applied the sensor to examine intracellular acid adaptation across diverse cancer cell systems. Although cancer cell lines are removed from their native tumor microenvironments during long-term culture, we reasoned that they may nevertheless retain a persistent physiological memory of adaptation to the chronically acidic conditions encountered in vivo. Building on our observation that acute extracellular acidification drives a corresponding reduction in intracellular pH (pHi), we reasoned that the extent to which cells preserve intracellular pH under acid stress could provide a quantitative readout of the retained adaptation state. We refer to this relationship between extracellular pH (pHe) and the resulting intracellular pH response as acid imprinting.

To systematically profile acid imprinting behavior, we transiently expressed cytoplasmic ORION in a panel of cancer and non-cancer cell lines spanning pancreatic cancer, melanoma, glioblastoma, and esophageal epithelial models (Fig. 4a). Cells were exposed to MES/HEPES-buffered PBS spanning extracellular pH 5.0–7.0, followed by ratiometric BRET measurements and conversion of sensor responses to intracellular pH values using purified ORION calibration curves. This workflow enabled direct quantitative comparison of intracellular buffering behavior across distinct cellular backgrounds in a scalable 384-well format.

**Figure 4.**
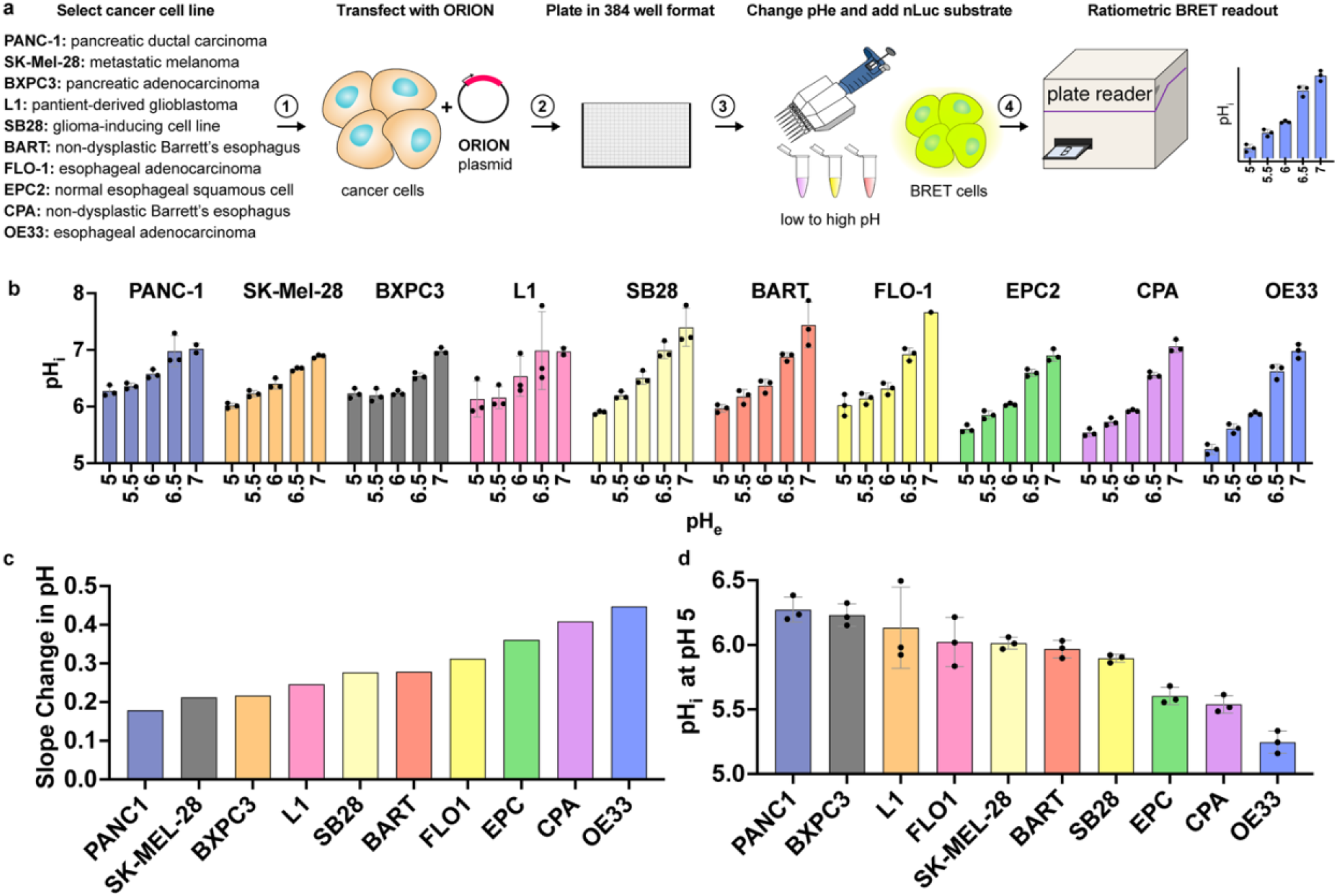
Characterizing acid imprinting across cancer cell models using ORION. (**a**) Schematic of the ORION-based acid imprinting workflow. Cancer and non-cancer cell lines were transiently transfected with cytoplasmic ORION, plated in 384-well format, and exposed to MES/HEPES-buffered PBS spanning extracellular pH (pH_e_) 5.0–7.0 in 0.5-unit increments. Following the addition of the NanoLuc substrate furimazine, ratiometric BRET measurements were acquired using a plate reader, and intracellular pH (pH_i_) values were estimated using purified ORION calibration curves. Cell lines examined included PANC-1 pancreatic ductal carcinoma, SK-MEL-28 metastatic melanoma, BxPC3 pancreatic adenocarcinoma, L1 patient-derived glioblastoma, SB28 glioma-inducing cells, BART non-dysplastic Barrett’s esophagus, FLO-1 esophageal adenocarcinoma, EPC2 normal esophageal squamous cells, CPA non-dysplastic Barrett’s esophagus, and OE33 esophageal adenocarcinoma cells. (**b**) Steady-state intracellular pH (pH_i_) measurements as a function of extracellular pH (pH_e_) across the indicated cell lines, revealing heterogeneous intracellular buffering behavior and acid adaptation profiles between cellular backgrounds. Data shown are representative of n = 3 biological replicates, with each biological replicate averaging 4 technical replicates per condition. (**c**) Calculated ΔpH_i_/ΔpH_e_ slopes for each cell line derived by linear regression across extracellular pH 5.0–7.0. The pH_i_-to-pH_e_ slope provides a quantitative measure of intracellular pH regulatory capacity and acid imprinting behavior across distinct cellular contexts. (**d**) Waterfall plot ranking cell lines by intracellular pH (pH_i_) following acute extracellular acidification to pH 5.0, highlighting substantial heterogeneity in acid resistance and maintenance of intracellular pH homeostasis across cancer and non-cancer cell models.

All cell lines exhibited progressive intracellular acidification in response to decreasing extracellular pH, but the magnitude of these responses varied substantially across models (Fig. 4b). Some cell types maintained relatively elevated intracellular pH despite strong extracellular acidification, whereas others exhibited marked intracellular acidification. These findings demonstrate substantial heterogeneity in acid-buffering capacity and suggest that distinct cancer-associated cellular states retain varying degrees of adaptation to acidic microenvironments.

To quantitatively compare these behaviors, we calculated ΔpHi/ΔpHe slopes across extracellular pH 5.0–7.0 for each cell line (Fig. 4c). Shallower slopes corresponded to stronger intracellular pH homeostasis and greater resistance to extracellular acidification, whereas steeper slopes reflected increased susceptibility to intracellular acidification. Notably, substantial variability was observed across cancer models, indicating that acid imprinting is not simply associated with transformation status but instead reflects heterogeneous physiological adaptation programs across cellular contexts.

We next ranked cell lines according to intracellular pH following acute extracellular acidification to pHe 5.0 (Fig. 4d). This analysis revealed pronounced differences in acid resistance across the panel. PANC-1 and BxPC3 pancreatic cancer cells maintained comparatively elevated intracellular pH under acidic stress, whereas OE33 esophageal adenocarcinoma cells exhibited substantially reduced intracellular pH. Intermediate behaviors were observed across glioblastoma, melanoma, and Barrett’s esophagus-derived models. Together, these findings support the existence of heterogeneous acid imprinting states across cancer cell lineages and suggest that tumor cells can retain persistent physiological memory of adaptation to acidic microenvironments even after prolonged culture outside their native in vivo environments. Importantly, these observations were enabled by ORION’s expanded acidic range, which extends beyond that of pHluorin2 and allows quantitative resolution of intracellular acidification states that are poorly captured by existing fluorescent pH sensors.

## Discussion

Our findings reveal that pH responsiveness is an intrinsic and previously underappreciated property of widely used BRET systems. Although BRET architectures are commonly treated as spectroscopically stable platforms for measuring molecular interactions and signaling events, both luciferase donors and fluorescent acceptors exhibited substantial environmental sensitivity across physiologically relevant pH conditions. Importantly, these effects were amplified within intact donor–acceptor architectures, resulting in large shifts in spectral output under acidic conditions. These observations have important implications for the interpretation of BRET-based assays used throughout cell biology, pharmacology, and biosensor engineering. Cellular processes associated with altered proton homeostasis, including receptor trafficking, endosomal maturation, metabolic stress, hypoxia, and extracellular acidification, may directly influence BRET behavior independently of the biological phenomenon under study. Rather than representing isolated idiosyncrasies of specific proteins, our results suggest that environmental pH sensitivity is a broader systems-level property embedded within commonly used BRET architectures.

At the same time, these findings demonstrate that the intrinsic environmental responsiveness of BRET systems can be rationally repurposed for biosensor engineering. By exploiting rather than suppressing pH sensitivity, we developed ORION, a genetically encoded ratiometric BRET pH sensor optimized for acidic and physiological biological environments. ORION combines the advantages of excitation-free luminescence imaging with a broad operating range in acidic conditions, a large dynamic response window, and strong quantitative performance in living cells. These properties are particularly important because many biologically and clinically relevant compartments exist outside the optimal range of commonly used fluorescent pH sensors, including lysosomes, endosomes, metabolically stressed tissues, and acidic tumor microenvironments. The expanded acidic range of ORION further enabled quantitative resolution of intracellular acidification states that became compressed using pHluorin2. Collectively, these features position ORION among the most sensitive genetically encoded tools currently available for interrogating acidic intracellular environments in living systems.

Beyond biosensor development, our results also identify acid imprinting as a potentially important feature of cancer cell physiology. Although cultured cancer cell lines are removed from their native tumor microenvironments, distinct cellular backgrounds retained markedly different capacities to resist intracellular acidification during acute extracellular acid stress. These findings suggest that adaptation to acidic tumor environments may persist as a stable physiological state even after prolonged ex vivo culture^35^. The mechanisms underlying this phenomenon remain to be determined, but may involve durable alterations in proton transport systems, membrane composition, organelle organization, metabolic state, or transcriptional and epigenetic programs linked to acid adaptation^36,37^. More broadly, these observations raise the possibility that intracellular pH regulation itself may encode aspects of tumor evolutionary history and microenvironmental selection pressure. In this framework, acid imprinting may represent a measurable physiological memory of prior adaptation to acidic tumor ecosystems.

Looking forward, ORION should provide a broadly useful platform for studying pH dynamics across molecular, cellular, and organismal systems. The sensor is readily compatible with subcellular targeting strategies, longitudinal live-cell measurements, and potentially in vivo imaging applications where excitation-independent approaches offer substantial advantages. In parallel, the concept of acid imprinting opens new directions for investigating how tumor cells adapt to chronic metabolic and microenvironmental stress and whether these states contribute to invasion, therapeutic resistance, immune interactions, or metastatic competence. More broadly, our findings establish environmental responsiveness not only as an important consideration in interpreting BRET-based assays but also as a powerful design principle for engineering next-generation genetically encoded biosensors.

## Methods

### Sensor Design & Cloning

Genetically encoded BRET-based pH sensors were constructed by fusing NanoLuc (Gift from Nevin Lambert), RLuc8, or GFP2 Trupath (Addgene Kit #1000000163) to either mVenus (gift from Nevin Lambert) or GFP2 via a flexible GGSGGS linker. BRET pairs were selected for testing pH-responsive energy transfer properties based on their widespread use in GPCR biosensors and complementary emission profiles. Constructs were assembled using HiFi DNA Assembly (NEB #E2621) following PCR amplification of each component. Primers included linker sequences and overhangs compatible with the multiple cloning sites of target plasmids. For initial screening in yeast, assembled constructs were cloned into the pYEplac181 episomal vector under the control of the TEF1 promoter. Cloning reactions were transformed into *E. coli* DH5α for propagation, and inserts were sequence-verified by Sanger sequencing (Eurofins). Positive clones were transformed into *S. cerevisiae* BY4741 for high-throughput BRET screening. The top-performing construct, ORION, consisting of NanoLuc fused C-terminally to mVenus via a GGSGGS linker, was subcloned into pcDNA3.1(+) for mammalian expression.

### High-Throughput Yeast Screening

Each BRET construct was overexpressed in *Saccharomyces cerevisiae* (strain BY4741) using the episomal vector pYEplac181 under the TEF1 promoter. Yeast transformation was performed via the lithium acetate/PEG method, and transformants were selected on synthetic dropout media lacking leucine (SC–Leu). Individual colonies were picked and cultured overnight in SC–Leu with 2% glucose, then diluted 1:50 into the fresh media and grown at 30 °C with shaking to an optical density (A_600_) of ∼0.8. For cell permeabilization, 9 mL of each yeast culture was mixed with 1 mL of digitonin solution (1 mg/mL in PBS, pH 7.0; Sigma-Aldrich, #D141) in a 50-mL conical tube. Samples were vortexed briefly and incubated at 30 °C for 10 minutes with shaking. After permeabilization, cells were centrifuged at 3,000 × g for 3 minutes, washed with 25 mL of PBS (pH 7.0), and pelleted again. The resulting cell pellet was resuspended in 25 mL of fresh PBS (pH 7.0). To conduct pH titrations, 1 mL aliquots of the permeabilized yeast suspension were transferred into eight 1.5 mL microcentrifuge tubes. Cells were pelleted by centrifugation, cleared of supernatant, and each resuspended in 1 mL of pre-warmed pH-buffered solutions: PBS adjusted to pH 5.0, 5.5, 6.0, 6.5, 7.0, 7.5, and 8.0, or Tris-KCl buffer (25 mM Tris, 100 mM KCl, pH 8.5). Buffers were adjusted using HCl or NaOH and prepared fresh to ensure pH accuracy.

For spectral acquisition, 36 μL of each pH-conditioned cell suspension was transferred into white 384-well plates (Greiner Bio-One, #781095). To initiate the bioluminescent reaction, 4 μL of Nano-Glo substrate (Promega, #N1110) diluted to 1× working concentration in PBS was added to each well. Bioluminescence emission spectra were recorded using a ClarioStar plate reader at a gain of 2500, capturing emission from 400 to 600 nm at 2 nm intervals. Raw data were background-subtracted and normalized to the emission intensity at pH 5.0. BRET ratios were calculated by dividing the fluorescent protein peak (e.g., 530 nm for mVenus) by the luciferase peak (e.g., 450 nm for NanoLuc), and plotted against buffer pH to generate titration curves.

### Bacterial Expression & Protein Purification

For bacterial expression, ORION, NanoLuc, mVenus, and pHluorin2 were cloned into pET-22b(+) (NovoPro, #V011019) for His-tag purification into competent BL21(DE3) cells. Transformants were grown overnight in 5 mL Luria broth + carbenicillin (100 μg/mL; Sigma-Aldrich, #C1389). The next morning, the overnight culture was transferred into 800 mL autoinduction media (ZYM-5052; prepared from ZY base [tryptone, RPI #T60060-500.0; yeast extract, RPI #P20240-500.0], 50× 5052 [D-(+)-glucose, RPI #G32045-3000.0; glycerol, VWR #97062-452; α-lactose monohydrate, Sigma-Aldrich #L2643], 50× M [Na_2_HPO_4_, Sigma-Aldrich #795410; KH_2_PO_4_, Sigma-Aldrich #795488; NH_4_Cl, Sigma-Aldrich #A9434; Na_2_SO_4_, Sigma-Aldrich #239313], and MgSO_4_ stock [Sigma-Aldrich #746452]; Studier, 2005), grown at 37 °C with shaking (200 rpm) for 8 h, and then grown overnight at 18 °C with shaking (200 rpm). The 800-mL autoinduction culture was split over two 500-mL centrifuge bottles, and cells were harvested by centrifugation at 4,500 rpm for 30 min. The supernatant was removed, and the pellet was resuspended in 35 mL PBS-TCEP (25 mM KPO_4_, 100 mM KCl, 1 mM TCEP [TCI Chemicals, #T1656], pH 7.0), transferred to 50-mL tubes, and stored at −20 °C. To purify overexpressed ratiometric pHluorin, cell pellets were thawed on ice, supplemented with fresh TCEP (TCI Chemicals, #T1656; to 1 mM), and lysed using a NanoDeBEE homogenizer (BEE International) at 30,000 psi. Following lysis, 250 μL of 5% polyethylenimine (PEI; Sigma-Aldrich, #408727; pH 7.9) was added to precipitate genomic DNA. The cell lysate was transferred to a 50-mL tube and centrifuged at 15,000 rpm for 30 min. The supernatant was then transferred to a 15-mL tube and incubated with 1 mL His bead slurry (Sigma-Aldrich, #P6611) for 15 min at 4 °C. The beads were harvested by centrifugation at 1,000 rpm for 1 min and then washed three times with PBS-TCEP. The beads were then transferred to a 2-mL spin column (Thermo Scientific, #89896) and eluted with PBS-TCEP + 250 mM imidazole (Sigma-Aldrich, #I0250). The eluted protein was collected in a 15-mL tube by centrifugation at 1,000 rpm for 1 min, transferred to a slide-a-lyzer dialysis cassette (Thermo Scientific, #87730), and dialyzed overnight in 4 L PBS-TCEP at 4 °C. Purified pHluorin stocks were stored at −80 °C.

#### pHluorin calibration

A pHluorin standard curve was calculated using purified ratiometric pHluorin. 200 μL PBS buffer titrated to each pH (137 mM NaCl, 2.7 mM KCl, 10 mM Na_2_HPO_4_, 1.8 mM KH_2_PO_4_) supplemented with 20 mM MES (Sigma-Aldrich, #M3671) or 20 mM HEPES (Sigma-Aldrich, #H3375, adjusted to the indicated pH values using HCl or NaOH) were placed into a 96-well microplate (CytoOne; CC7626-7596), and 36 μL aliquots were moved in technical quadruplicate to a black bottom 384-well plate (Greiner, 781096). Briefly, 4 μL ratiometric pHluorin (1 μM) were resuspended in each pH PBS buffer. The plate was vortexed at 2000 rpm for 30 seconds before reading. Excitation spectra were collected using a ClarioStar microplate reader (BMG LabTech) with the following parameters: top read, 40 flashes/ well, excitation start: 350 to 10 nm, excitation end: 495 to 10 nm, 5 nm steps; emission: 520 to 10 nm; instrument gain: 1700; focal height 7 mm. Raw fluorescence values at 385 and 475 nm were used to calculate the pHluorin ratio (385/475 nm) for each technical replicate. A standard curve was built by plotting the pHluorin ratio as a function of pH and fit using a sigmoidal 4PL model (X = log[concentration]) in GraphPad Prism. Experimental error was calculated in GraphPad Prism as the SD of the mean for the n = 4 technical replicates.

#### Orion calibration

An Orion standard curve was calculated using purified ratiometric nluc-mVenus. 200 μL PBS buffer titrated to each pH (137 mM NaCl, 2.7 mM KCl, 10 mM Na_2_HPO_4_, 1.8 mM KH_2_PO_4_) supplemented with 20 mM MES (Sigma-Aldrich, #M3671) or 20 mM HEPES (Sigma-Aldrich, #H3375, adjusted to the indicated pH values using HCl or NaOH) were placed into a 96-well microplate (CytoOne; CC7626-7596), ratiometric Orion (200nM final) were resuspended into each pH PBS buffer and 30 μL aliquots were moved in technical quadruplicate to a white plate 384-well plate (VWR, 82050-076) with white BackSeal (PerkinElmer, 6005199). Substrate Furimazine (Promega, N1110) was diluted into each pH of PBS, and 15μl Furimazine (0.75x) was added to 30μl of Orion for a final volume of 45μl (final Furimazine of 0.25x). The plate was vortexed at 1500 rpm for 30 seconds before reading. For full-spectrum luminescence, scans were collected using a ClarioStar microplate reader (BMG LabTech) with the following parameters: top read, measurement read time 1 second, emission start: 400 to 10 nm, emission end: 600 to 10 nm, 5 nm steps; instrument gain 3600, focal height 9.0 mm. To make the most optimal curve, endpoint scans were collected using a ClarioStar microplate reader (BMG LabTech) with the following parameters: top read, well multichromatics, chromatics were read at; first Chromatic: 450 to 10 nm; second chromatic 530 to 10 nm; instrument gain: 3600 for each chromatic; focal height 9.0 mm. A standard curve was built by plotting the Orion ratio as a function of pH and fit (sigmoidal, 4PL, X is log[concentration]) using GraphPad Prism. Experimental error was calculated in GraphPad Prism as the SD of the mean for the n = 4 technical replicates.

#### mVenus calibration

A mVenus reading was performed using purified mVenus. 200 μL PBS buffer titrated to each pH (137 mM NaCl, 2.7 mM KCl, 10 mM Na_2_HPO_4_, 1.8 mM KH_2_PO_4_) supplemented with 20 mM MES (Sigma-Aldrich, #M3671) or 20 mM HEPES (Sigma-Aldrich, #H3375, adjusted to the indicated pH values using HCl or NaOH) were placed into a 96-well microplate (CytoOne; CC7626-7596), and 36 μL aliquots were moved in technical quadruplicate to a black bottom 384-well plate (Greiner, 781096). Briefly, 4 μL mVenus (10 nM) were resuspended in each pH PBS buffer. The plate was vortexed at 2000 rpm for 30 seconds before reading. Excitation spectra were collected using a ClarioStar microplate reader (BMG LabTech) with the following parameters: top read, 40 flashes/ well, excitation start: 430 to 10 nm, excitation end: 510 to 10 nm, 5 nm steps; emission: 531.5to 21 nm; instrument gain: 2500; focal height 7 mm. Experimental error was calculated in GraphPad Prism as the SD of the mean for the n = 4 technical replicates.

#### Nanoluc calibration

A Nanoluc reading was performed using purified Nanoluc. 200 μL PBS buffer titrated to each pH (137 mM NaCl, 2.7 mM KCl, 10 mM Na_2_HPO_4_, 1.8 mM KH_2_PO_4_) supplemented with 20 mM MES (Sigma-Aldrich, #M3671) or 20 mM HEPES (Sigma-Aldrich, #H3375, adjusted to the indicated pH values using HCl or NaOH) were placed into a 96-well microplate (CytoOne; CC7626-7596), and 36 μL aliquots were moved in technical quadruplicate to a white plate 384-well plate (VWR, 82050-076) with white BackSeal (PerkinElmer, 6005199). Substrate Furimazine (Promega, N1110) was diluted into each pH of PBS, and 15μl Furimazine (0.75x) was added to 30μl of Nanoluc for a final volume of 45μl (final Furimazine of 0.25x). The plate was vortexed at 2000 rpm for 30 seconds before reading. For full-spectrum luminescence, scans were collected using a ClarioStar microplate reader (BMG LabTech) with the following parameters: top read, measurement read time 1 second, emission start: 410 to 10 nm, emission end: 530 to 10 nm, 5 nm steps; instrument gain 2000, focal height 11 mm. Experimental error was calculated in GraphPad Prism as the SD of the mean for the n = 4 technical replicates.

### Mammalian Cell Culture and Transfection

Human cell lines were obtained from ATCC, collaborators, or existing stocks and cultured under recommended conditions. Cancer lines included: PANC-1, SK-MEL-28, BxPC3 (ATCC); FLO-1, OE33, and non-cancer or pre-malignant lines included EPC2, BART, CPA(gift from El-Rifai); SB28 (murine glioma), and L1 (patient-derived GBM xenograft) (gift from Defne Bayik). PANC-1, SK-MEL-28, BxPC3, FLO-1, and L1 cells were cultured in DMEM (Thermo Fisher Scientific, Cat# 11995065) with 10% FBS (HyClone, #SH30071.03) and 1% Penicillin-Streptomycin-Glutamine (PSG, Cat# 10378016). OE33 and SB28 cells were maintained in RPMI-1640 (Thermo Fisher Scientific, Cat# 11875-119), also supplemented with 10% FBS and 1% PSG. EPC2, BART, and CPA cells were cultured in Complete Human Epithelial Cell Medium with Supplement Kit (Cell Biologics, Cat# H6621) according to the supplier’s instructions. All cells were incubated at 37 °C with 5% CO_2_ and passaged using TrypLE Express (Thermo Fisher, Cat# 12604013). For transient expression experiments, cells were seeded in 6-well plates and transfected using TransIT (Mirus Bio #MIR 2305). After 24 h, cells were trypsinized, counted, and replated into white 384-well plates (Greiner Bio-One #781095) at 5,000–10,000 cells per well for downstream assays.

### Acid Stress Assay and Phenotypic Classification

For acid stress assays, each cell line was seeded into the aforementioned white Greiner 384-well plates at a density of 5,000–10,000 cells per well and allowed to adhere for 48 h post-transfection or post-transduction. Cells were washed with pre-warmed PBS and incubated in pH-adjusted buffer (137 mM NaCl, 2.7 mM KCl, 10 mM Na_2_HPO_4_, 1.8 mM KH_2_PO_4_) supplemented with 20 mM MES (Sigma-Aldrich, #M3671) or 20 mM HEPES (Sigma-Aldrich, #H3375, adjusted to the indicated pH values using HCl or NaOH)) for 10 minutes at room temperature. pH values ranged from 5.0 to 7.0 in 0.5-unit increments. Each well received 30 μL of buffer at the indicated pH.

To avoid buffer dilution artifacts, furimazine (Nano-Glo, Promega, cat# N1110) was pre-diluted in the same pH buffer and added at a 0.25x final concentration to a volume of 15 μL per well, bringing the total volume to 45 μL. Plates were sealed and incubated at room temperature in the dark for a total of 10 minutes following buffer addition before luminescence acquisition. Bioluminescence was measured using a ClarioStar plate reader with the following parameters: top read, well multichromatics, chromatics were read at; first Chromatic: 450 to 10 nm; second chromatic 530 to 10 nm; instrument gain: 3600 for each chromatic; focal height 9.0 mm. The BRET ratio was calculated as the acceptor emission divided by the donor emission. Three biological replicates were performed for each condition.

## Author contributions

D.G.I. managed the study, and D.G.I. and S.T. wrote the manuscript. S.T., K.L., B.C., J.A., and S.C performed all experiments.

## Acknowledgements

This work was supported by the National Institutes of Health through an R35 Maximizing Investigators’ Research Award (R35GM119518 to D.G.I.), which provided core funding for this study. Additional support was provided by a Pap Corps Champions for Cancer Research Endowed Chair to D.G.I. and the Sylvester Comprehensive Cancer Center.

## References

1 Isom, D. G. et al. Protons as second messenger regulators of G protein signaling. Mol Cell 51, 531–538 (2013). 10.1016/j.molcel.2013.07.012

2 Webb, B. A. et al. Structures of human phosphofructokinase-1 and atomic basis of cancer-associated mutations. Nature 523, 111–114 (2015). 10.1038/nature14405

3 Andersson, M. et al. Proton-coupled dynamics in lactose permease. Structure 20, 1893–1904 (2012). 10.1016/j.str.2012.08.021

4 Webb, B. A., Chimenti, M., Jacobson, M. P. & Barber, D. L. Dysregulated pH: a perfect storm for cancer progression. Nat Rev Cancer 11, 671–677 (2011). 10.1038/nrc3110

5 Swietach, P. What is pH regulation, and why do cancer cells need it? Cancer Metastasis Rev 38, 5–15 (2019). 10.1007/s10555-018-09778-x

6 Morales Rodriguez, L. M., Crilly, S. E., Rowe, J. B., Isom, D. G. & Puthenveedu, M. A. Location-biased activation of the proton-sensor GPR65 is uncoupled from receptor trafficking. Proc Natl Acad Sci U S A 120, e2302823120 (2023). 10.1073/pnas.2302823120

7 Kapolka, N. J. et al. Proton-gated coincidence detection is a common feature of GPCR signaling. Proc Natl Acad Sci U S A 118 (2021). 10.1073/pnas.2100171118

8 Rowe, J. B., Kapolka, N. J., Taghon, G. J., Morgan, W. M. & Isom, D. G. The evolution and mechanism of GPCR proton sensing. J Biol Chem 296, 100167 (2021). 10.1074/jbc.RA120.016352

9 Kapolka, N. J. et al. DCyFIR: a high-throughput CRISPR platform for multiplexed G protein-coupled receptor profiling and ligand discovery. Proc Natl Acad Sci U S A 117, 13117–13126 (2020). 10.1073/pnas.2000430117

10 Casey, J. R., Grinstein, S. & Orlowski, J. Sensors and regulators of intracellular pH. Nat Rev Mol Cell Biol 11, 50–61 (2010). 10.1038/nrm2820

11 Nishi, T. & Forgac, M. The vacuolar (H+)-ATPases--nature’s most versatile proton pumps. Nat Rev Mol Cell Biol 3, 94–103 (2002). 10.1038/nrm729

12 Ballabio, A. & Bonifacino, J. S. Lysosomes as dynamic regulators of cell and organismal homeostasis. Nat Rev Mol Cell Biol 21, 101–118 (2020). 10.1038/s41580-019-0185-4

13 Piasentin, N., Milotti, E. & Chignola, R. The control of acidity in tumor cells: a biophysical model. Sci Rep 10, 13613 (2020). 10.1038/s41598-020-70396-1

14 Diwu, Z., Chen, C. S., Zhang, C., Klaubert, D. H. & Haugland, R. P. A novel acidotropic pH indicator and its potential application in labeling acidic organelles of live cells. Chem Biol 6, 411–418 (1999). 10.1016/s1074-5521(99)80059-3

15 Chen, X. et al. pH sensing controls tissue inflammation by modulating cellular metabolism and endo-lysosomal function of immune cells. Nat Immunol 23, 1063–1075 (2022). 10.1038/s41590-022-01231-0

16 Larkin, J. R., Foo, L. S., Sutherland, B. A., Khrapitchev, A. & Tee, Y. K. Magnetic Resonance pH Imaging in Stroke - Combining the Old With the New. Front Physiol 12, 793741 (2021). 10.3389/fphys.2021.793741

17 Michl, J., Park, K. C. & Swietach, P. Evidence-based guidelines for controlling pH in mammalian live-cell culture systems. Commun Biol 2, 144 (2019). 10.1038/s42003-019-0393-7

18 Mahon, M. J. pHluorin2: an enhanced, ratiometric, pH-sensitive green florescent protein. Adv Biosci Biotechnol 2, 132–137 (2011). 10.4236/abb.2011.23021

19 Reifenrath, M. & Boles, E. A superfolder variant of pH-sensitive pHluorin for in vivo pH measurements in the endoplasmic reticulum. Sci Rep 8, 11985 (2018). 10.1038/s41598-018-30367-z

20 Shen, Y., Rosendale, M., Campbell, R. E. & Perrais, D. pHuji, a pH-sensitive red fluorescent protein for imaging of exo-and endocytosis. J Cell Biol 207, 419–432 (2014). 10.1083/jcb.201404107

21 Tantama, M., Hung, Y. P. & Yellen, G. Imaging intracellular pH in live cells with a genetically encoded red fluorescent protein sensor. J Am Chem Soc 133, 10034–10037 (2011). 10.1021/ja202902d

22 Rennick, J. J., Nowell, C. J., Pouton, C. W. & Johnston, A. P. R. Resolving subcellular pH with a quantitative fluorescent lifetime biosensor. Nat Commun 13, 6023 (2022). 10.1038/s41467-022-33348-z

23 Surana, S., Bhat, J. M., Koushika, S. P. & Krishnan, Y. An autonomous DNA nanomachine maps spatiotemporal pH changes in a multicellular living organism. Nat Commun 2, 340 (2011). 10.1038/ncomms1340

24 Yue, X. et al. DNA-Based pH Nanosensor with Adjustable FRET Responses to Track Lysosomes and pH Fluctuations. Anal Chem 93, 7250–7257 (2021). 10.1021/acs.analchem.1c00436

25 Goyet, E., Bouquier, N., Ollendorff, V. & Perroy, J. Fast and high resolution single-cell BRET imaging. Sci Rep 6, 28231 (2016). 10.1038/srep28231

26 Salahpour, A. et al. BRET biosensors to study GPCR biology, pharmacology, and signal transduction. Front Endocrinol (Lausanne) 3, 105 (2012). 10.3389/fendo.2012.00105

27 Yang, J. et al. Coupling optogenetic stimulation with NanoLuc-based luminescence (BRET) Ca(++) sensing. Nat Commun 7, 13268 (2016). 10.1038/ncomms13268

28 Pfleger, K. D. & Eidne, K. A. Illuminating insights into protein-protein interactions using bioluminescence resonance energy transfer (BRET). Nat Methods 3, 165–174 (2006). 10.1038/nmeth841

29 Jang, W., Lu, S., Xu, X., Wu, G. & Lambert, N. A. The role of G protein conformation in receptor-G protein selectivity. Nat Chem Biol 19, 687–694 (2023). 10.1038/s41589-022-01231-z

30 Namkung, Y. et al. Monitoring G protein-coupled receptor and beta-arrestin trafficking in live cells using enhanced bystander BRET. Nat Commun 7, 12178 (2016). 10.1038/ncomms12178

31 Olsen, R. H. J. et al. TRUPATH, an open-source biosensor platform for interrogating the GPCR transducerome. Nat Chem Biol 16, 841–849 (2020). 10.1038/s41589-020-0535-8

32 Wan, Q. et al. Mini G protein probes for active G protein-coupled receptors (GPCRs) in live cells. J Biol Chem 293, 7466–7473 (2018). 10.1074/jbc.RA118.001975

33 Rowe, J. B., Lee, K. & Isom, D. G. PIONEER: A periplasmic display platform for synthetic biology-based screening of genetically encoded protein regulators. J Biol Chem 302, 110967 (2026). 10.1016/j.jbc.2025.110967

34 Rowe, J. B., Taghon, G. J., Kapolka, N. J., Morgan, W. M. & Isom, D. G. CRISPR-addressable yeast strains with applications in human G protein-coupled receptor profiling and synthetic biology. J Biol Chem 295, 8262–8271 (2020). 10.1074/jbc.RA120.013066

35 Sutoo, S., Maeda, T., Suzuki, A. & Kato, Y. Adaptation to chronic acidic extracellular pH elicits a sustained increase in lung cancer cell invasion and metastasis. Clin Exp Metastasis 37, 133–144 (2020). 10.1007/s10585-019-09990-1

36 Michl, J. et al. Acid-adapted cancer cells alkalinize their cytoplasm by degrading the acid-loading membrane transporter anion exchanger 2, SLC4A2. Cell Rep 42, 112601 (2023). 10.1016/j.celrep.2023.112601

37 Ordway, B., Swietach, P., Gillies, R. J. & Damaghi, M. Causes and Consequences of Variable Tumor Cell Metabolism on Heritable Modifications and Tumor Evolution. Front Oncol 10, 373 (2020). 10.3389/fonc.2020.00373

